# Network-enabled efficient image restoration for 3D microscopy of turbid biological specimens

**DOI:** 10.1101/2020.05.27.118869

**Authors:** Le Xiao, Chunyu Fang, Yarong Wang, Tingting Yu, Yuxuan Zhao, Dan Zhu, Peng Fei

## Abstract

Though three-dimensional (3D) fluorescence microscopy has been an essential tool for modern life science research, the light scattering by biological specimens fundamentally prevents its more widespread applications in live imaging. We hereby report a deep-learning approach, termed ScatNet, that enables reversion of 3D fluorescence microscopy from high-resolution targets to low-quality, light-scattered measurements, thereby allowing restoration for a single blurred and light-scattered 3D image of deep tissue, with achieving improved resolution and signal-to-noise ratio. Our approach can computationally extend the imaging depth for current 3D fluorescence microscopes, without the addition of complicated optics. Combining ScatNet approach with cutting-edge light-sheet fluorescence microscopy, we demonstrate that the image restoration of cell nuclei in the deep layer of live Drosophila melanogaster embryos at single-cell resolution. Applying our approach to two-photon excitation microscopy, we could improve the signal and resolution of neurons in mouse brain beyond the photon ballistic region.

## Introduction

A recurring challenge in biology is the attempt to look inside various biological specimens to study the structure, function of their cellular constituents [1, 2]. These increasing quests are posing substantial challenges to the current 3D light microscopy. Conventional 3D optical microscopy methods based on linear one-photon absorption processes, no matter epiillumination [3–6] or plane illumination modes [7–10], are limited to the imaging of the tissue surface (less than 100 μm) because highly anisotropic scattering blurs the images at increasingly greater depths. Tissue clearing methods [11–14] generally use chemical reagent to remove the scattering cellular constituents and then select high refractive index solution for refractive index matching, thereby suppressing the multiple light scattering and substantially increasing the imaging depth. However, tissue clearing can’t completely eliminate the scattering effect, especially for very thick disordered medium [15–17]. More importantly, cleaning methods requires tissue extraction and histological preparation of the sample, making it impossible for imaging intact tissue or living organisms [14]. Nonlinear two-photon-excited fluorescence microscopes (TPEM) provide superior optical sectioning while mitigate the light scattering effect, owing to the dominance of ballistic photons inherent in the nonlinear two-photon laser excitation process and the use of longer excitation wavelength [18]. However, the contribution of ballistic fluorescence emission photons remains minor beyond about one scattering mean-free-path (50–90 μm at 600 nm in brain gray matter), and becomes negligible several hundred microns deep into brain tissue [18–20]. Temporal focusing TPEM with generalized phase contrast [16] could further increase the imaging depth to a few hundreds of microns, but at the cost of more acquisition time and thereby being difficult for live imaging.

Aside from the direct acquisition of scattering-suppressed image, several hardware-based computational methods, such as adaptive optics [21, 22], and speckle correlation [23] techniques, have been proposed to recover scattering images by extracting useful information from the blurred signals. To compensate for the wavefront distortion of the incident beam, adaptive optics utilizes wavefront sensing components in conjunction with the guide star feedback mechanism to optimize the incident light as well as the fluorescence emission in-and-out of the turbid medium. Thus, it relies on highly sophisticated optics as well as extra response time of wavefront measurement and correction, making it less compatible with commercialized confocal or TPE microscopes, and less suited for observing highly dynamic samples. Speckle correlation methods utilize the correlation of laser speckle imaging and optical memory effects, but the angular range of optical memory effects limit the imaging depth below the superficial layers of tissues.

Instead of recovering scattered signals based on conventional optical models, the recently-emerging deep neural networks can be another promising alternative, which can learn end-to-end image mapping relationship from data pairs without the need of explicit analytical modeling [24]. Deep learning based on convolutional neural networks (CNNs) [15] has recently emerged in the fields of enhancing biomedical images quality. For example, fluorescence microscopy has recently benefited from the advances in deep-learning-based enhancement such as image restoration [25], deconvolution [26, 27], super-resolution [28–31], and style transformation [32–34]. Also, deep-learning has been newly applied to the suppression of light scattering, but limited to the demonstration of 2D photography [35].

Here we propose deep-learning-enable computation pipeline, termed ScatNet, which can efficiently restore blurred signals induced by tissue scattering in a 3D fluorescence image stack with the enhancement of spatial resolution and image contrast. Based on multi-view acquisition and registration by light-sheet microscopy, and stepwise tissue thickening experiment by TPEM, we construct a faithful 3D dataset containing well-registered scattering and scattering-free image pairs for ScatNet (U-Net architecture) training. Afterward, a well-trained ScatNet is then capable of directly predicting a deblurred high-quality image from the scattering input, without requiring any hardware addition or complicated computation. To demonstrate its enhancement to various microscopy modalities, we applied ScatNet restoration to the single-view SPIM image of Drosophila embryo, and successfully recovered the completely blurred nuclei structures at the deep region of embryo, thereby showing its potential to substitute more complicated multi-view SPIM imaging strategy. We also demonstrate ScatNet can double the penetration depth for commercialized TPEM imaging of tagged neurons in Thy1-YFP mouse brain tissues.

## Results

### ScatNet for tissue scattering suppression

We acquire well-registered pairs of 3D images which show different degree of light scattering by our delicately designed setups using multi-view light-sheet microscopy or two-photon excitation microscopy. Two pairs of acquired raw images are first normalized and cropped into many patches to fit the network training (**Fig. 1a, step 1**). A background filter is then applied to reduce their backgrounds (**Fig. 1a, step 2**). After data pre-processing, the network corresponds these low-quality scattering and high-quality scattering-free images as target data and label data, respectively, to initiate the training process (**Fig. 1b, step 3**). During each epoch of the training, the neural network basically learns how to better restore high-quality images from the scattering target inputs. The generated intermediate output by each epoch is compared with the fixed label data to optimize the loss function of the network, and thereby iteratively push the network toward its optimized stage, at which the network can predict high-quality outputs close enough to the label data (**Fig. 1b, step 4, 5, 6**). After the network being efficiently trained, then apply it to the restoration of 3D images acquired from biological imaging experiment, e.g., live imaging, which is often vulnerable to light scattering (**Fig. 1c, step 7**). By computationally removing the blurs by light scattering, ScatNet significantly improves the signal-to-noise ratio (SNR) and resolution of 3D images (**Fig. 1d, step 8, 9, 10**). In the following sections, we demonstrate how ScatNet can optimize the LSFM and TPEM imaging in live imaging of biological specimens, with substantially increasing the imaging depth.

**Fig. 1.**
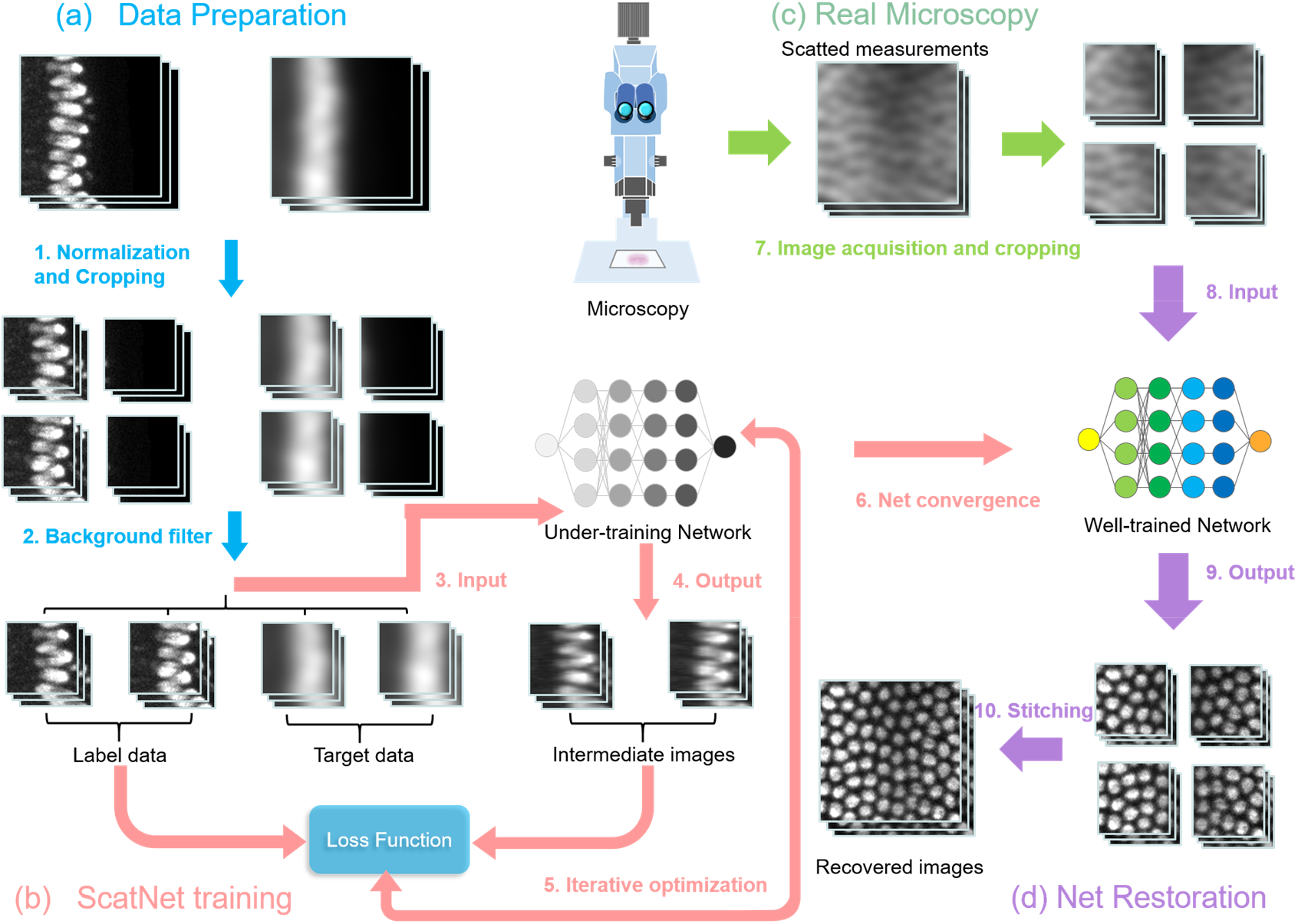
Workflow of ScatNet. **(a)** Data pre-preparation including the intensity normalization, size cropping (step 1), and background filtration (step 2) of the raw 3D image pairs acquired by imaging setups that exclude/include effect of light scattering. **(b)** Iterative network (U-Net architecture) training based on the registered and preprocessed image pairs. The network generates intermediate outputs based on the input target data (scattering images) and compares them quantitatively with the label data (scattering-free images), to calculate the system loss function which can iteratively push the optimization of the network (step 3 to 6). **(c)** Real microscopy imaging of deep tissues (Drosophila embryo, mouse brain, etc.), which generates experimental images with low SNR and resolution by light scattering (step 7). **(d)** Restoration of the degraded experimental images through predicting higher-resolution, higher-SNR images (step 8, 9) using the well-trained ScatNet. The restored patches are finally stitched back into a complete 3D image volume of the sample (step 10).

### Characterization of ScatNet

We first characterized the performance of ScatNet through the point-spread-function (PSF) imaging of sub-diffraction fluorescent beads (~500 nm in diameter, Lumisphere) using a Bessel light-sheet microscope built on an upright microscope (Olympus BX51, 800-nm light-sheet thickness, 20×/0.5 detection objective). The beads were embedded in an agarose column (1%), mixed with a cluster of cultured human cervical cancer (Hela) cells (~500 × 500 × 600 μm3), serving as a scattering medium to provide light scattering for PSF imaging. In the experiment, we rotated the sample 180° to flip it upside down, thereby able to image the same beads twice with and without cells’ scattering medium coupled at 0°, scattering mode, and 180°, scattering-free mode, respectively. After imaging some regions of interest, a 3D image registration was applied to precisely align the scattering and scattering-free PSF images for network training. Then we imaged another region of beads in the same procedure, to acquire the scattered image and corresponding ground truth for network application (**Fig. 2a**). The ScatNet has successfully restored low-quality scattering image. The result show that, spatial resolution improved obviously as well as high similarity to ground-truth image (**Fig. 2b**). The magnified views of the ***yz*** and ***xz*** projections of a selected region of interest (ROI) further reveal the details of the imaged/restored beads. The linecuts through the individual beads at measurement (***xy***) and reconstructed (***xz***) planes confirmed much narrower full-width-at-half-maximums (FWHMs) of restored PSFs, as compared to the original ones (**Fig. 2c, d**). With referring to the scattering-free ground truth, the peak signal to noise ratio (PSNR) and structure similarity (SSIM) values of network-restored signals are improved across the imaging depth [36, 37] (**Fig. 2e, f**). In addition to the fluorescent beads, we further recovered a negative USAF resolution target (Thorlabs R3L3S1N) imaged through a 300-μm thickness chicken breast slice (**Supplementary Fig. S2**). The restoration of the completed blurred and distorted line pairs also validated the performance of ScatNet.

**Fig. 2.**
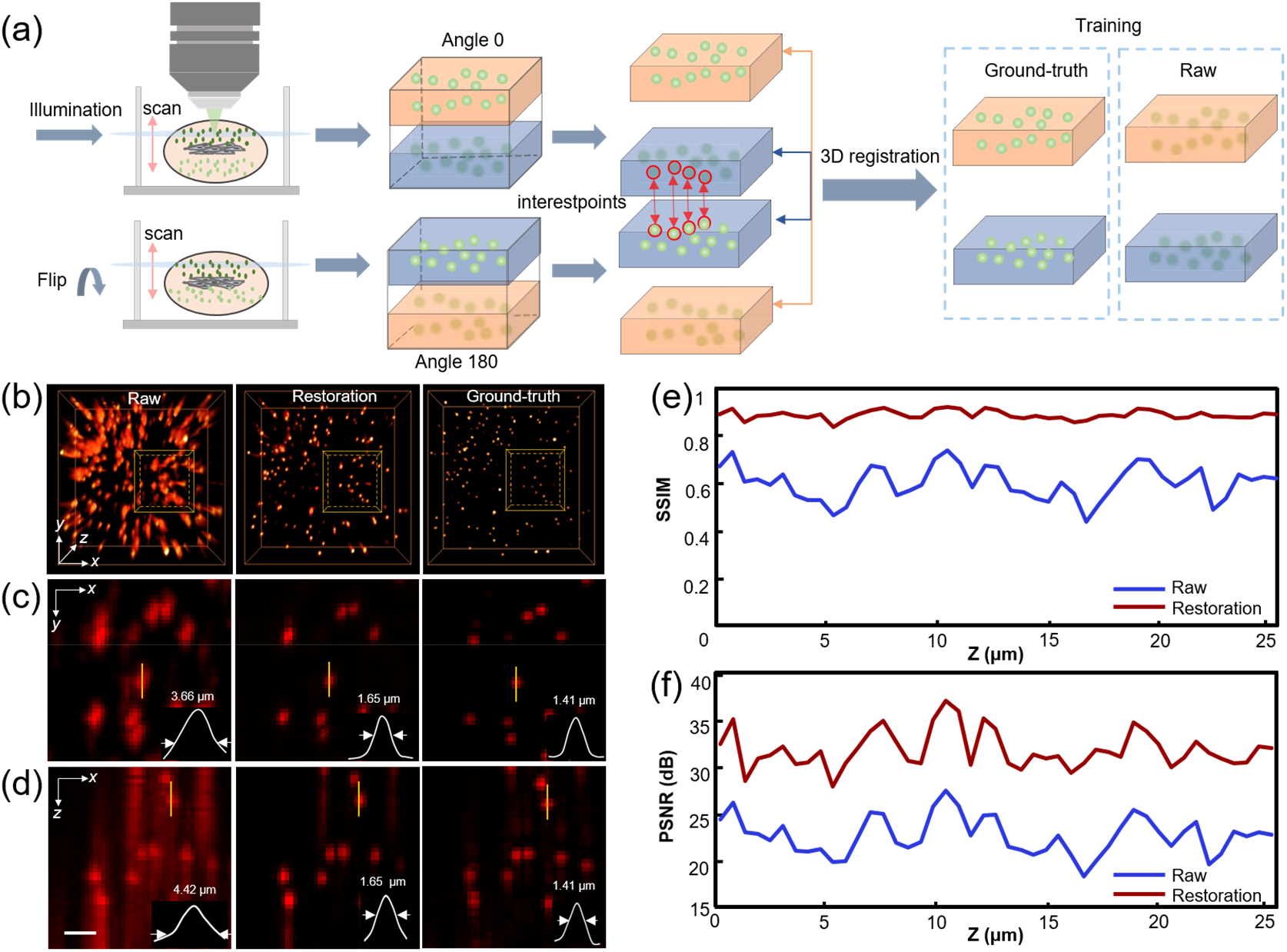
Performance of ScatNet. **(a)** Light-sheet microscopy imaging of 3D fluorescent beads buried with a cluster of scattering Hela cells. Two image stacks of the same beads are acquired under two opposite views (0° and 180°), and spatially registered to obtain pairs of scattering and scattering-free bead images. **(b)** 3D reconstructions of the same beads by scattering mode (raw input), scattering-free mode (ground truth), and ScatNet restoration (network output). **(c)-(d)** Axial (***xy***) and lateral (***xz***) maximum intensity projections (MIPs) of the same selected ROIs (yellow boxes in **b**), comparing the spatial resolutions (insets, FWHMs of PSFs) by the three modes. The results clearly demonstrate a notable improvement in lateral and axial resolution for ScatNet. ScatNet improves the lateral/axial resolution across the entire ~25-μm overlapping volume, with achieving relatively uniform resolution of ~1.6 ± 0.1 μm and ~1.6 ± 0.2 μm in ***x/y*** and ***z***, respectively, which are compared to ~3.6 ± 0.1 μm and ~4.4 ± 0.2 μm in the raw scattering image. Scale bar, 5 μm. **(e)** The structure similarity (SSIM) of the ScatNet restoration as compared with the scattering-free ground truth image. The results show sufficiently high restoration fidelity by ScatNet. **(f)** The layer-by-layer peak signal to noise ratio (PSNR) of the ScatNet restoration, with using GT as reference.

### Improvement for light-sheet microscopy of live Drosophila melanogaster embryos

We applied ScatNet to the SPIM imaging of live Drosophila melanogaster embryos, which suffers from strongly tissue scattering majorly from the lipids of the embryo body. Multi-view LSFM imaging as we mentioned above were thereby used to obtain a set of LSFM stacks (8-16 groups), which are further registered and fused to reconstruct a 3D image containing complete sample signals. In our implementation, we first used open SPIM images of Drosophila embryos for ScatNet training [36]. We divided 7-view data into three groups with each containing a pair of corresponding upside-down 3D images (0°-180°, 45°-225°, 90°-270°). Following the above-mentioned registration and preprocessing steps, three groups of training data contain well-registered scattering/scattering-free signals at deep/superficial layers for the network training to make neural network predictions, which could directly recover the blurred low-contrast nuclei signals of single-view SPIM image (**Fig. 3b, c**). Thereby, our network recovery procedure eliminated the use of multi-view acquisition as well as computation-demanding registration/fusion. Three selected ROIs that suffer from mild, medium, and strong scatterings were restored by our ScatNet and compared with their corresponding ground truths (**Fig. 3d**). The network predictions are verified to have sufficiently high structural similarity (**Fig. 3d**), improved resolution (**Fig. 3e**), and enhanced signal-to-noise ratio (SNR, **Fig. 3f**), which are adaptable to all the regions regardless of the degree of tissue scattering.

**Fig. 3.**
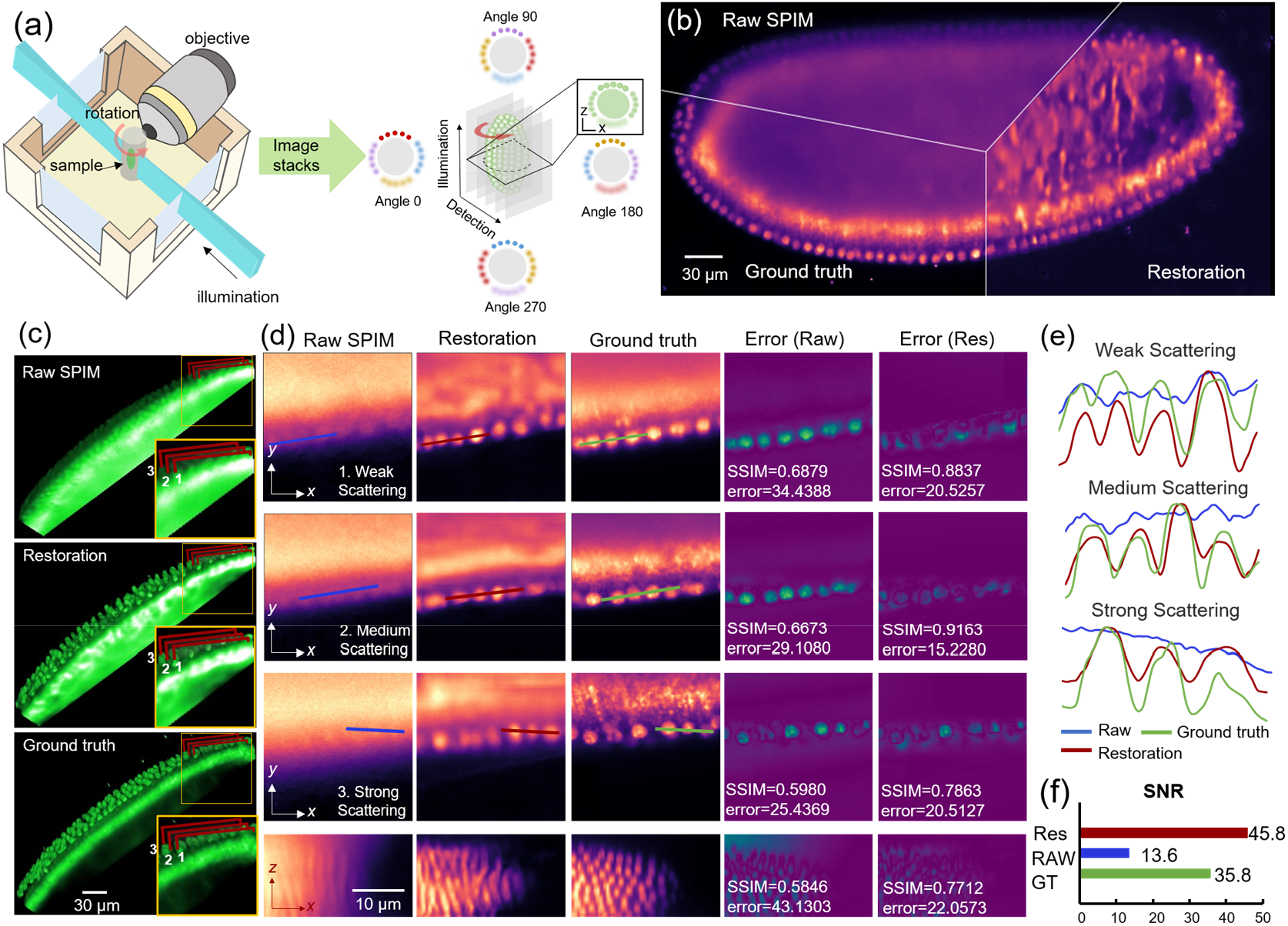
Restoration of living Drosophila embryos imaged by LSFM. **(a)** 3D image stacks of Drosophila embryos acquired by Multi-view SPIM and registered by multi-view registration [38]. The 3D stacks from different views are registered to obtain the training pairs of scattering and scattering-free images from different parts of the embryo. **(b)** The reconstructed 3D Drosophila embryo by raw SPIM (scattered signals), multi-view fusion (ground-truths) and ScatNet (restored signals). **(c)** 3D views of raw, restored and ground-truth signals of nuclei at the same embryo region. **(d)** ScatNet restoration of signals with different degrees of light scattering (at different depths). The comparative results are shown in both raw ***x-y*** (row 1-3) and reconstructed ***x-z*** planes (row 4). The error maps of raw and restored images, as compared to ground truths, are correspondingly shown in the right two columns. **(e)** Intensity plots of the lines through the nuclei resolved in raw, restored, and ground-truth images. The plot profiles of ScatNet fit those of ground-truths well, showing sufficiently high resolution as well as restoration fidelity have been achieved regardless of the scattering degree. **(f)** The averaged SNR values of the image stacks shown in **(c)**. It is obvious that the poor SNR in raw images has been also improved by ScatNet.

### Improvement for two-photon excitation microscopy of mouse brain

Light microscopy imaging of live model animals, such as mouse and rat, has become a type of widely-used technique for life science research. Though optical imaging can provide highly specific structural and functional information, the observation is often limited to very superficial layer owing to the light scattering in tissue. For example, for in vivo two-photon excitation microscopy, the ballistic regime of photons (~920 nm) in a mouse brain tissue is usually limited to ~100 μm in depth, which merely corresponds to the Cortex I area in an intact brain. To restore optimal imaging performance at depth, various methods have been proposed to increase the imaging depth of current light microscopy, but most of them requiring the adaptive optics or other extra hardware to be added into the original microscopes.

Here we used ScatNet to computationally solve the scattering problem and substantially increase the imaging depth for a commercialized two-photon excitation microscope (Olympus FV1000, **Fig. 4a**), without the need of hardware retrofit. To quantitatively mimic the scattering effect at different imaging depth, we first imaged (Olympus XLPLN, 25×/1.05 water dipping objective) the YFP-tagged neurons in a coronal slice of mouse brain (100-μm thickness, Thy1-YFP), then we imaged the same brain slice shielded by a series of homologous brain tissue slices with different thickness of 100, 200 and 300 μm (**Fig. 4a**). Accordingly, we acquired four groups of 3D image stacks of neurons, serving as the ground-truth data (no shield, signals depth 0-100 μm), weak scattering data (100-μm shield, signals depth 100-200 μm), medium scattering (200-μm shield, signals depth 200-300 μm), and strong scattering data (300-μm shield, signals depth 300-400 μm). Since we carefully stabilized the sample slice when adding or removing the shield slices, the acquired four groups of 3D images were naturally aligned. In our ScatNet implementation, we paired the three groups of scattering images with the ground-truth images to construct a training system containing hierarchical scattering models (**Fig. 4a**). Then we applied the well-trained ScatNet to the blurred images of neurons acquired at different scattering levels, i.e. with different shield depths (**Fig. 4b3, b5, b7**), and compared the results with raw images (**Fig. 4b2, b4, b6**), as well as the ground truth (**Fig. 4b1**). We noticed that the restored images show notably more details of neuron fibers compared to the ambiguous scattering inputs. The linecuts through individual neuronal fibers (**Fig. 4b1 to b7**) resolved at 100 to 300-μm depth further confirmed narrower FWHMs by ScatNet restoration (plots in **Fig. 4c**). Refer to the scattering-free ground truth, we also compared the PSNR and SSIM values of the raw scattering and network-restored images across the entire imaging depth **(Fig. 4d, e**). These quantitative analyses have verified the capability of our ScatNet to enhance the two-photon microscopy image quality for deep tissues imaging. Besides the above-mentioned validation experiment, we also successfully demonstrated the application of ScatNet to the signal recovery of a 300-μm turbid brain tissue (**Supplementary Fig. S3**).

**Fig. 4.**
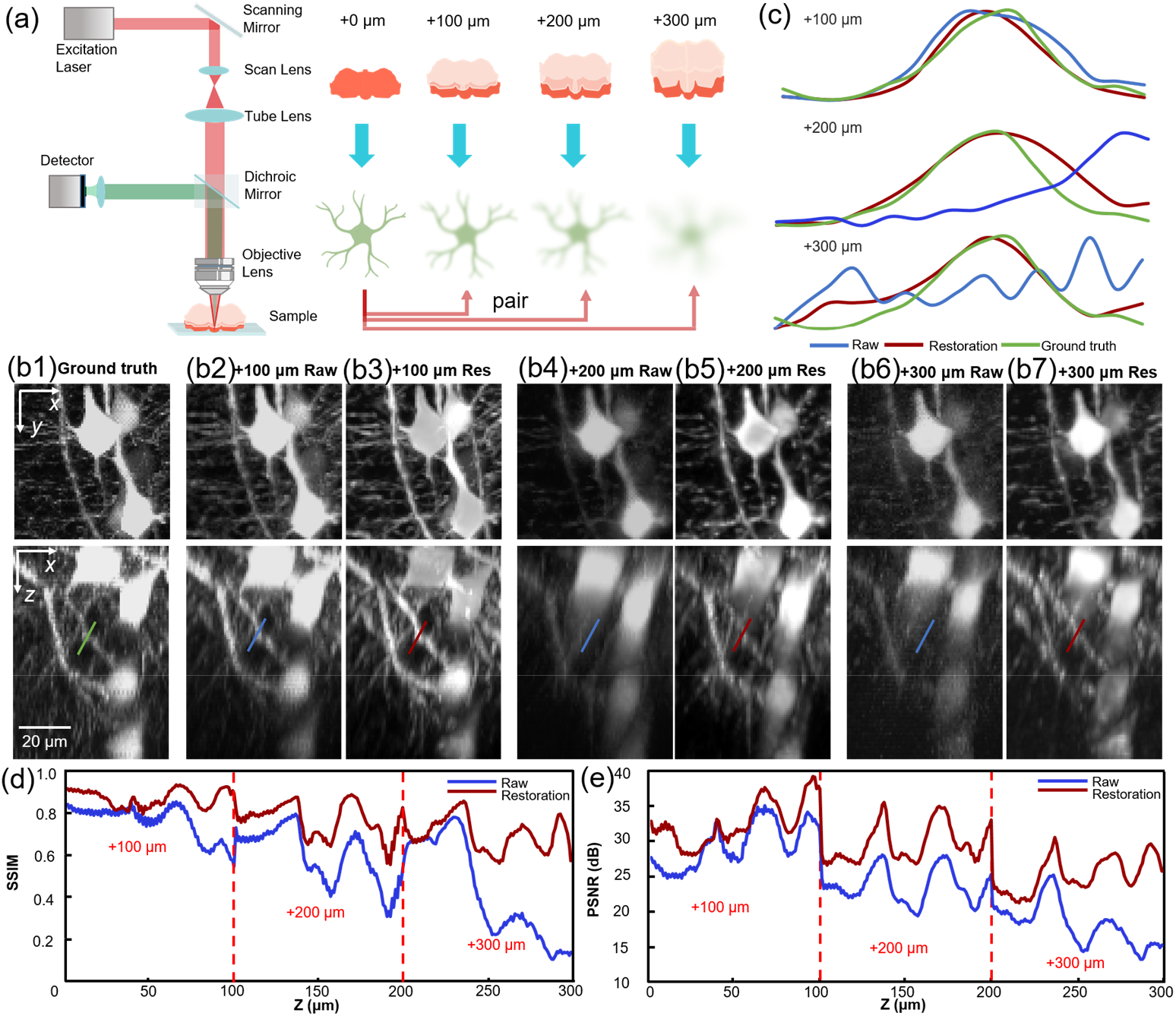
ScatNet restoration of neuronal signals in mouse brain slices (Thy1-YFP-M) imaged by TPEM. **(a)** The experimental design of TPEM imaging for obtaining the gradient scattering information through covering the sample with different thickness of brain slices. The network trained by data encompassing gradient scattering information is then able to recover signals suffering from different degrees of scattering. **(b)** Maximum intensity projections (MIPs) of the same 70 × 70 × 100 μm^3^ regions along ***z***- (top) and ***y***-axis (bottom). **(b1)(b2)(b4)(b6)** show the raw imaging results through 0 (ground-truth), 100, 200, and 300 μm brain slices, respectively. **(b3)(b5)(b7)** correspondingly show the results restored from the scattered signals in **(b2)**, **(b4)**, and **(b6)**, respectively. **(c)** Intensity plots of linecuts through individual neuronal fibers resolved in **b1** to **b7**. The comparative intensity profiles quantitatively verify improved resolution as well as high restoration fidelity by ScatNet for different degrees of scattering. **(d)(e)** Variation of SSIM and PSNR values of the raw measurements and corresponding ScatNet restorations, across the imaging depth. The ScatNet improvement is more obvious at deeper region.

## Discussion

Unlike classical approaches introducing extra sophisticated optics or algorithm to suppress light scattering, our data-driven method is purely computational, based on powerful prediction capability of deep neural network. In addition, ScatNet contributes to simple, fast image restoration without laborious manual operations. In our ScatNet applications, we significantly restored the scattering-induced blurring for light-sheet microscopy imaging of live Drosophila embryos, thereby allowing in-vivo observation of embryo development with a simpler lightsheet microscope setup. We also demonstrated that ScatNet substantially increase imaging depth of intact mouse brain for two photon excitation microscopy, without involving adaptive optics. The performance of ScatNet restoration, including the achieved resolution and signal accuracy, were also quantitatively analyzed and verified to be far better than the raw scattering inputs. Despite these advantages, as a proof-of-concept, our network method remains highly data dependent, relying on the precise registration of experimental scattering and scattering-free data, which are difficult to obtain in some applications. We envision this can be possibly improved by recent advances in Generative Adversarial Network (GAN) which can correlate unpaired images at the superficial and deep side, or the development of image degradation algorithm that can generate intrinsically aligned synthetic scattering images from those scattering-free measurements. Also, our method still has to compromise between the higher recovery fidelity and stronger scattering-induced degradation. We expect this could be addressed with the unceasing development of deep learning, allowing more versatile architectures and powerful training pipelines.

In summary, we have proposed a deep-learning-based method that can restore 3D microscopy image degradation by light scattering in biological tissues. The versatile enhancements of our method in wide-field, light-sheet, and two-photon excitation microscopes have been sufficiently verified via imaging various types of biological tissues. Therefore, our method provides a paradigm for readily addressing the challenge of light scattering, which widely exists in deep tissue imaging. Furthermore, through computationally reconstruction of a 3D image with drastically improved quality, more in-depth biological applications, e.g., imaging beyond the superficial cortex area, or more accurate image-based analyses, e.g., region segmentation/annotation, could possibly be realized in a simpler and more efficient way.

## Funding sources and acknowledgements

National Natural Science Foundation of China (21874052 for P.F., 61860206009 for D.Z.), National Key R&D program of China (2017YFA0700501, D.Z. and P.F.), Innovation Fund of WNLO (P.F.) and Junior Thousand Talents Program of China (P.F.). We thank Hao Zhang for the help on the code implementation.

See Supplement 1 for supporting content.

## Notes

### Competing Interest Statement

The authors have declared no competing interest.

