## Supplementary figures and notes for "Network-enabled efficient image restoration for 3D microscopy of turbid biological specimens"

### 1. Deep network structure

Restoration of scattered images is an inverse problem aiming to reconstruct a clear output image from a degraded input. In this work, we directly obtain the nonlinear mapping function between input-output using deep neural network rather than those complicated optical scattering models. Herein, a U-Net structure was used as it has achieved good performance in different biomedical applications. Due to the elastic deformation data enhancement, U-Net only requires a small amount of label images and relatively shorter training time [1, 2].

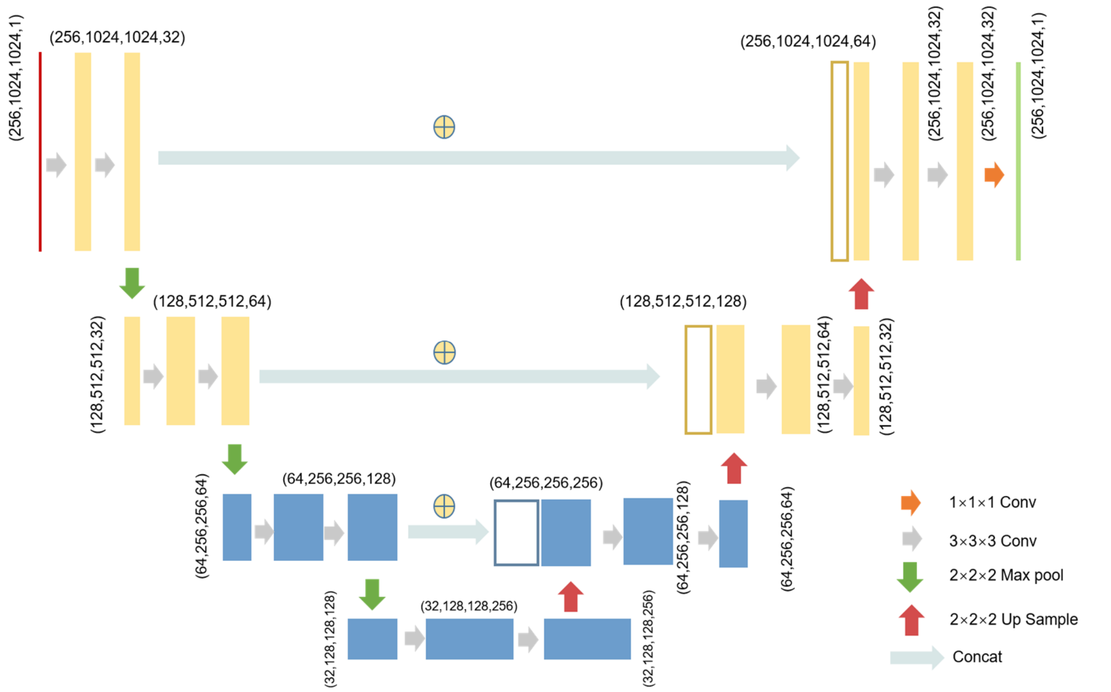


**Fig. S1. Illustration of our U-Net-based network.** This learning procedure contains an encoder-decoder architecture with skip-connections. The "Conv" represents a convolutional layer. The 3×3×3 Conv layers use the Rectified Linear Unit (ReLU) as the active function. "Max pool" is the MaxPooling3D layer. "Up Sample" is the UpSampling3D layer. "Concat" means concatenation operation.

### 2. ScatNet on USAF resolution target with wide-field microscope

Before demonstrating the capability of ScatNet for recovering 3D image of biological sample, we first verified our approach on the 2D image of a standard USAF resolution target (Thorlabs R3L3S1N). A wide-field microscope (Olympus BX51, 4×/0.1 objective) was used to image the resolution board with/without the coverage of a layer of tissue phantom (frosted tape), for obtaining the light-scattered and corresponding scattering-free image pairs, respectively. with highest resolution of 102.0 line pairs per mm (lpm). Then we used ~80% amount of such automatically aligned image pairs for ScatNet training, and the rest ~20% data for validation. Before network implementation, the training images were cropped into 128×128-pixel sizes and a background filter was applied to filtrate the informative images. An image rotation was applied to these pre-processed images to further augment the data scale to 3750 image pairs. After the network training achieving convergence, we validated its performance on the testing data (another 20%), which are not included in the network before. The results showed that our ScatNet is capable of restoring the line pairs (32.0 to 102.0 lines per mm, lpm), which were all blurred and distorted in the fuzzy raw images, owing to the strong and non-uniform light scattering by the tissue phantom (**Fig. S2**).

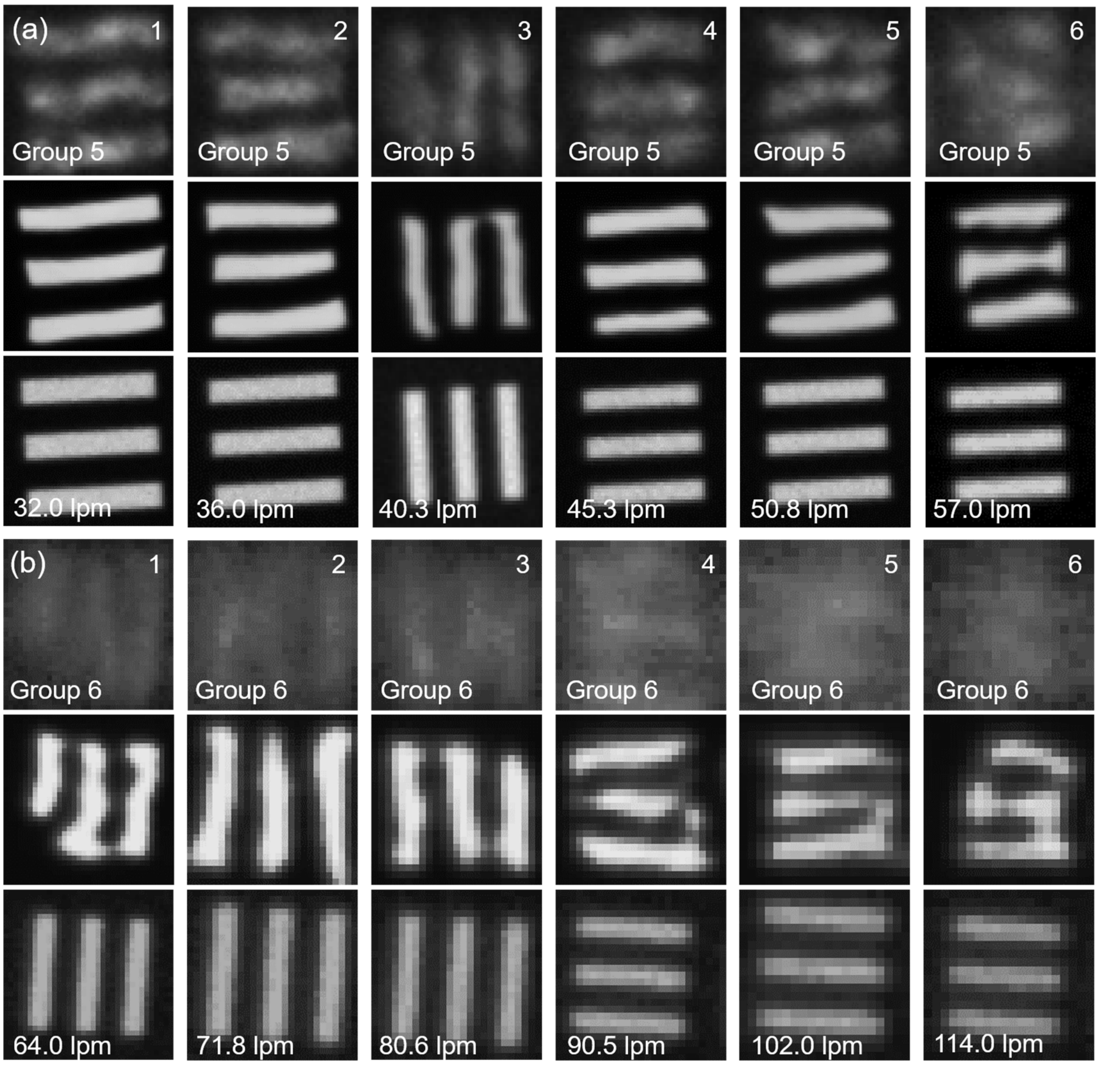


**Fig. S2. ScatNet restoration of 2D image of USAF resolution target acquired through a layer of tissue phantom.** Raw (tissue phantom covered), ScatNet-restoration and ground-truth (tape removed) images of the same area in the USAF resolution target, shown as top, middle, and bottom parts, respectively. **(a)** Comparative results of single line pairs in group 5 (32.0 to 57.0 lpm) before and after network restoration. **(b)** The highest resolution of single line pairs (group 6, 64.0 to 114.0 lpm) can be restored by our ScatNet.

#### 3. Drosophila embryo

We downloaded the dataset containing 7-view SPIM acquisitions of Drosophila embryo from http://fly.mpi-cbg.de/preibisch/nm/HisYFP-SPIM.zip. Among those 3D stacks, we used the bead-based registration algorithm to collect low- and high scattering images from overlapping regions obtained from three opposite views for network training and testing. The tutorial on how to use the registr-ation plugin in basic and advanced mode is available at http://pacific.mpi-cbg.de/wiki/index.php/SPIM_Registration.

### 4. Brain slices imaging

Before imaging, the sample brain slice (neurons labelled by Thy1-YFP) was mounted onto a glass slide substrate. Then the packed sample was carefully fixed on the microscope stage with avoiding the shift caused by adding/removing scattering tissue slices. We used a commercialized TPE microscope (Olympus FV 1000, Olympus XLPLN, 25×/1.05 water dipping objective) to image neurons in sample brain slices beneath different depths of scattering tissues. To obtain clear 3D volumes for network training, we first directly imaged the 100 μm-thickness sample brain slices (size of each frame 512×512 pixels, step size 2 μm, 1×1×2 $\mu m^3$ voxel). The imaging parameters for obtaining ground-truth data were: 920-nm pulse excitation with 10% power and 0-10% gain, corresponding to a power of 3-5 mW under the objective. Then, to obtain light-scattered fuzzy 3D images for network training as well as validation, we added 1 to 3 blank brain slices (100 μm-thickness each, no fluorescence labelling) onto the sample slice step by step, to simulate the tissue scattering at different depths from 0 to 400 μm. As the imaging depth of sample signals became larger with thicker blank tissues added, the laser power was also gradually increased to obtain signals with enough SNR for subpixel registration with the ground truths. Finally, after 300 μm-thickness blank tissue added, the neuron signals became too weak to be processed for network training, indicating a maximum imaging depth of ~400 μm in our experiment.

### 5. ScatNet restoration for TPEM image of a 300-μm brain slice

Beside the performance validation, we also practically applied our trained network to the 3D TPEM image of a 300-μm thick brain slice. The restoration results are shown and compared with raw inputs in **Fig. S3**. We specifically compared planes at depths of 60, 180, and 240 μm, which represented superficial, middle, and deep layers in raw and restored image stack to demonstrate our model's high robustness to different scattering situations (**Fig.S3(c)**). It is clearly revealed from both ***xy*** measurement planes and ***xz*** reconstructed planes that our ScatNet restoration is able to maintain the original high-quality signals at the superficial layer and at the same time, recover degraded signals at deep layer with providing increased resolution and contrast. The SNR values for different depths are improved both visually and quantitatively.

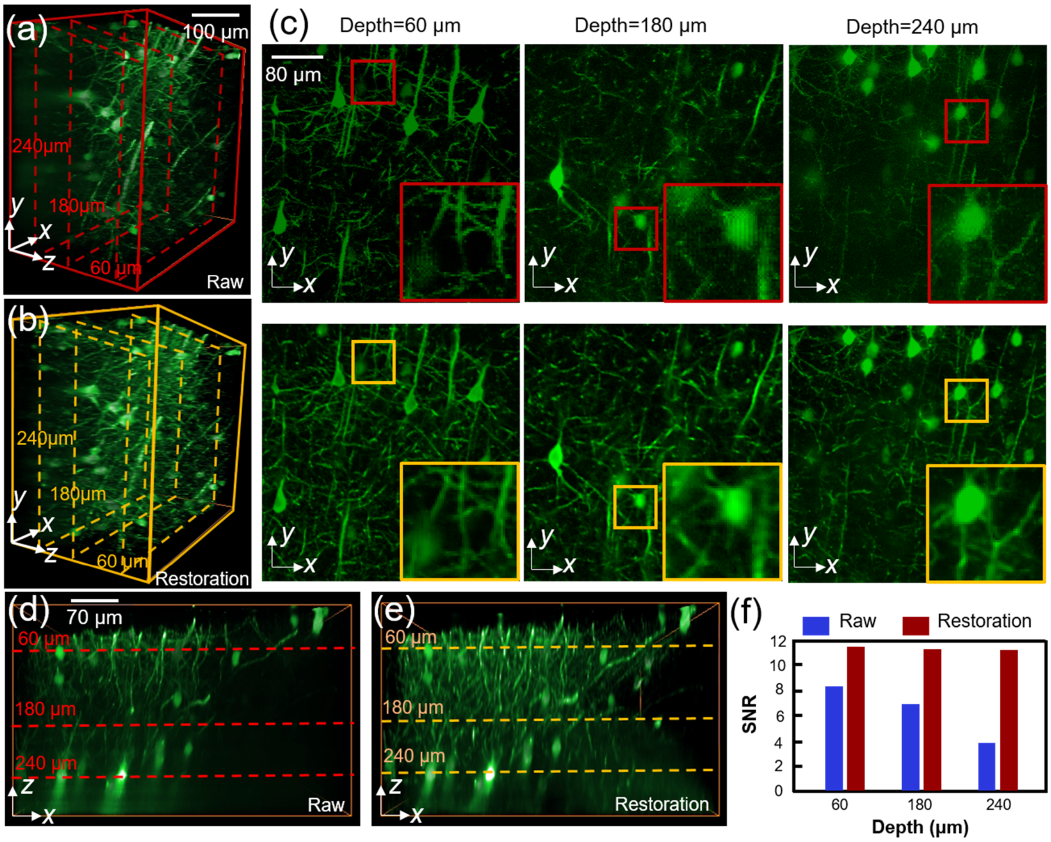


**Fig. S3. Restoration of 300-μm thick brain slice with neurons labelled by YFP (Thy1-YFP-M). (a), (b)** 3D views of the raw and restored neurons in the same brain volume across entire 300-μm depth. Scale bar, 100 μm. **(c)** Axial (***xy***) planes of the selected ROIs (dashed boxes in **a** and **b**) at 60 μm, 180 μm and 240 μm depth in the raw (upper row) and restoration (lower row) results, respectively. Scale bar, 80 μm. **(d), (e)** The ***xz*** reconstruction planes in the raw and ScatNet-restored 3D images, showing significantly enhanced neuron details with improved contrast at the deep side. Scale bar, 70 μm. **(f)** Comparison of SNR values at abovementioned three different depths.

### 6. Comparison of several network implementation methods

We also note that the network parameters, including architectures, and loss functions, could also have effects on the final restored results. Here, we selected a region of interests (ROI) ~80 × 80 × 100 $\mu m^3$ in the 200-μm shield 100-μm thick Thy1-YFP-M brain slice to validate the accuracy and generalization capability of a few different deep-learning models. With using the same loss function (Mean Square Error, MSE or Cross Entropy, CE), we compared the performance of Resnet with our U-Net-based ScatNet on the same brain neuron dataset, as shown in **Fig.S4(c)-(f)**. This difference is possibly because the influence of long and short skip connections [3]. On the other hand, with using the same network architecture, either U-Net-based ScatNet or ResNet, we found that MSE might not be suitable for handling deep tissue scattering tasks and loss function of CE performs better (**Fig.S4(c), (d) vs (e), (f)**). Compared to pure denoising tasks, deep tissue scattering is more complex owing to the multiple deflection of lights randomly distributed across the whole image. In this case, CE loss aims to find the global distribution with maximum likelihood, which is more effective than conventional MSE [4] (**Fig.S4(f)**).

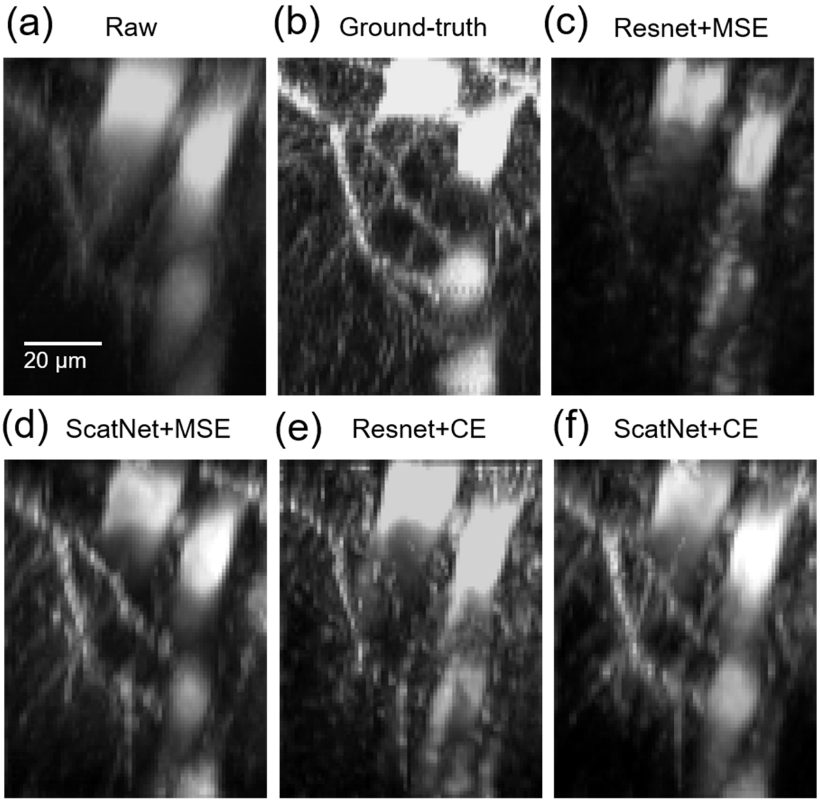


**Fig. S4. Comparison of the restored results by different deep-learning models with different parameters. (a)-(f)** lateral (*xz*) maximum intensity projections (MIPs) of the same selected ROIs of raw scattering image, ground-truth scattering-free image, restoration by ResNet with MSE loss function, restoration by ScatNet with MSE loss function, restoration by ResNet with CE loss function, and restoration by ScatNet with CE loss function, respectively. Scale bar, 20 µm.
